# Pre-symptomatic reduction of individuality in the App^NL-F^ knock-in model of Alzheimer’s disease

**DOI:** 10.1101/2022.08.18.504396

**Authors:** Fanny Ehret, Meike S. Pelz, Anna N. Senko, Karla E. G. Soto, Hang Liu, Gerd Kempermann

## Abstract

While one third of the risk for Alzheimer’s disease (AD) is explained by environment and lifestyle, AD pathology also affects individual lifestyle, possibly long before clinical manifestation of dementia. To study this hidden disease effect with its potentially large impact on coping with AD, we examined in mice how the App^NL-F/NL-F^ (NL-F) knock-in mutation affects the pre-symptomatic response to environmental enrichment (ENR). We assessed the emergence of inter-individual phenotypic variation while both genetic background and the shared environment were held constant, thereby isolating the contribution of individual behavior (‘non-shared environment’). After 4 months of ENR, in NL-F both mean and variability of plasma ApoE were increased, suggesting a pre-symptomatic variation in pathogenic processes. Roaming entropy (RE) as a measure of behavioral activity was continuously assessed with radiofrequency identification (RFID) technology and revealed reduced habituation and variance in NL-F compared to controls. Intra-individual variation decreased but behavioral predictability and stability were reduced in NL-F. Seven months after discontinuation of ENR we found no difference in plaque size and number, but ENR increased variance in hippocampal plaque counts in NL-F. A reactive increase in adult hippocampal neurogenesis in NL-F, known from other models, was normalized by ENR. Our data suggest that, while NL-F has early effects on individual behavioral patterns in response to ENR, there are lasting effects at the level of cellular plasticity even after discontinuation of ENR. Hence, early behavior matters for maintaining individual behavioral trajectories and brain plasticity even under maximally constraint conditions.

## Introduction

While in large epidemiological studies the genetic risk to develop Alzheimer’s disease has been estimated to be two thirds, the remaining risk can be attributed to modifiable risk factors (1). Targeting these factors for prevention offers a powerful, feasible and economic way to reduce the danger that AD cases will overwhelm the health care systems worldwide within the coming few decades. Particularly counteracting physical inactivity and low educational attainment have a very high potential for prevention (2–4). The reality of beneficial lifestyles, however, is much more complex, subtle and individual than the prominently recommended interventions with the greatest effect size suggest (5, 6). In addition, the timing of lifestyle intervention is crucial, given that administration during symptomatic stages might be too late for successful intervention (7, 8). Beneficial lifestyles would have to be adopted early during pre-clinical stages in order to exert their maximum effect.

To better understand the emergence of individual responses to the progressing disease and the relationship between behavioral trajectories and individual resilience, we made use of a unique experimental paradigm that exposes the behavioral (non-shared) component of the environmental factor in phenotypic variation (9, 10), when genes are kept constant (11). We have previously found that in the hippocampus, such individualizing effects on behavior and adult neurogenesis can be lasting, even if external stimuli are discontinued, and are associated with epigenetic changes at genes involved in plasticity (12, 13).

Impaired adult neurogenesis is increasingly discussed as a relevant contributing mechanism underlying memory deficits in AD, although some controversy exists in respect to adult neurogenesis as a compensatory mechanism and its link to plaque pathology and Braak stage (14–17). Because adult neurogenesis is a strongly individualizing trait, but by itself is affected by transgenic over-expression of AD-related genes, which might obscure the interaction effect with behavior. We therefore used a 2^nd^ generation mouse model of AD, in which artificial overexpression is no longer a confound, due to the knock-in of the disease mutation into the physiologically regulated mouse gene (8). Young App knock-in mice can be considered as models of preclinical AD and therefore have an ideal genetic background to study the influence of environmental factors. The App^NL-F/NL-F^ knock-in mice carry the Swedish [NL] and Beyreuther/Iberian [F] mutation in the amyloid-precursor protein (App) leading to elevated levels of amyloid-ß (Aß) 42 and increased ratio of Aß42/Aß40 (18). App^NL-F/NL-F^ knock-in mice (short: NL-F) slowly start to develop Aß plaques and gliosis by 6 months of age and the first cognitive impairments with 18 months (18) and thus are suitable for intervention studies before disease manifestation. App^NL/NL^ (short: NL) mice only have the Swedish mutation, which has no impact on the disease phenotypes (18).

Beginning at the age of 6 weeks, both NL and NL-F strains of mice were kept in the large ENR enclosure, which consists of 70 connected standard cages (Fig. 1B). They were constantly tracked with RFID transponders injected under their skin in the neck. After 4 months the mice were returned to their home cage for mid and late adulthood in order to mimic human conditions of increasing inactivity with advancing age as it is often seen in western lifestyle. The brains were analyzed at 13 months, after all mice had been tested behaviorally at the end of the 4 months ENR phase (Fig. 1A).

**Figure 1.**
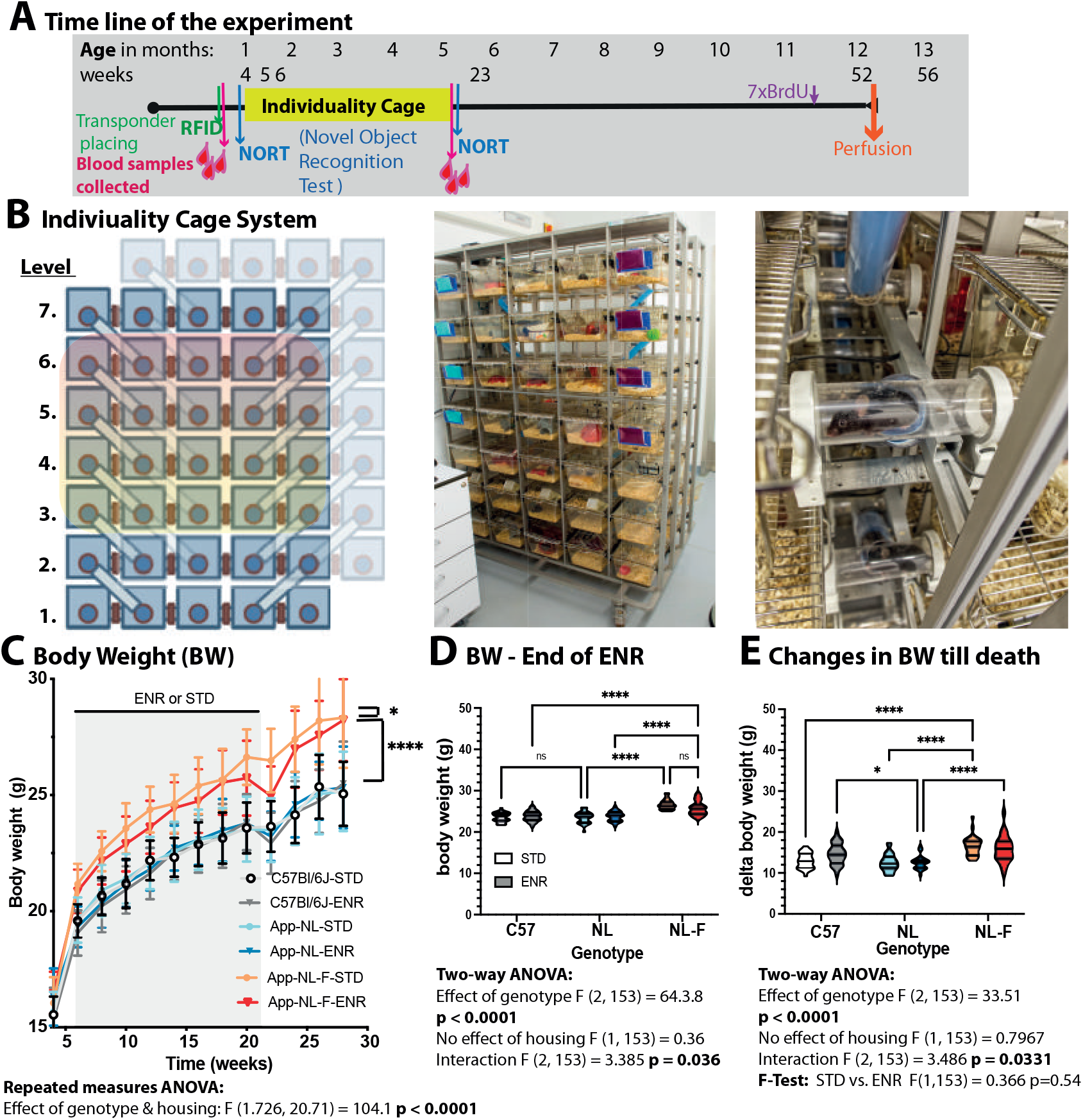
Experimental layout and body weight development of mice living in a complex enclosure. (**A**) Experimental time line of the experiment. (**B**) Schematic representation (left) and pictures (right) of the individuality cage system for complex enrichment (ENR) and longitudinal monitoring. Each of the 70 cages within the 7 levels contain RFID ring antennas (red) around the tunnels, thereby connecting the different levels and cages next to each other. Space and complexity increased over time from 3 to 7 levels. (**C**) App-NL-F (NL-F) mice show elevated body mass already prior to ENR which reduce upon ENR housing. (**D**) Body weight at the end of ENR at 6 months of age. (**E**) The gained body weight (BW) between 5 weeks till 13 months of age does don’t differ between control mice in STD or ENR condition, whereas NL-F animals increase body weight more pronounced, nevertheless variance is not significantly altered but a pronounced trend was seen as shown by F-test below. Performed statistical analysis are shown below each graph. Significant interactions are stated as ns = not significant, *p<0.05, **p<0.01, ***p<0.005.

With this experimental design we set out to examine the emergence of differences in individual behavioral trajectories in mice with AD predisposition (NL-F) vs. controls (NL), as such differences might represent an access to mechanisms, why pre-clinical stages of AD might already be associated with behaviors that support disease development or, conversely, prevent it. We thus also asked whether such behavioral differences might have a measurable effect on Aß pathology and the neurogenic reserve in the hippocampus (19). Besides the longitudinal parameters on behavioral activity, metabolic changes in blood, object discrimination and endpoint parameters of adult neurogenesis and pathology were analyzed.

## Material and Methods

### Animal husbandry

App^NL-F/NL-F^ and App^NL/NL^ mice were obtained from the Riken Institute and contain a Swedish (KM670/671NL) and Beyreuther/Iberian (I716F) mutation, as reported (18). C57Bl6J mice were used a second control line for the analysis of potential side effects, however no abnormalities were observed. A total of 162 female animals from the 3 genotypes were maintained on a 12 h light/dark cycle with food and water provided *ad libitum* at the Center for Regenerative Therapies (CRTD), Dresden, Germany. Control and enriched animals received the same fortified chow (no. V1534; Sniff) with 9% of energy from fat, 24% from protein, and 67% from carbohydrates. The experiment was licensed from the local authority (Regierungspräsidium Dresden) under DD24-5131/354/51. At the age of 4 weeks, half of the animals were randomly assigned to the two different housing conditions either living in custom-built system of enrichment (PhenoSys GmbH, now marketed as “PhenoSys ColonyRack Individuality 3.0”) referred to here as enriched housing (ENR) or standard housing (STD). The experimental groups consisted of 27 female animals per condition and genotype, housing 81 animals together in the same enriched environment and 81 animals in STD separated for each genotype. At 5 weeks of age, animals were randomly separated into the 2 condition and ENR were subcutaneously injected into their neck with a glass-coated microtransponders (SID 102/A/2; Euro I.D.) under brief isoflurane anesthesia. At the same time, blood was collected from the retrobulbar venous plexus from all animals. At 6 weeks, mice entered into the different housing conditions. For ENR the condition changed over time since initially mice used only 3 out of 7 levels from this cage-rack system (Fig. 1A). Every 2 weeks, mice were taken out of the system, a new level was made accessible, toys changed in position and complexity and cages and tunnels were cleaned. After 9 weeks, all 7 levels were available and only the position and diversity of toys, nesting material, and huts were changed. Food and water locations remained the same over the entire period. After 17 weeks of ENR, all mice left the cage system and were returned to home cages of 4-5 animals for the rest of the experiment until 13 months of age (Fig. 1D). At the time of exit, blood samples were again drawn from the venous plexus. In addition, body weight was monitored biweekly and glucose levels were measured every 4 weeks in a drop of blood from the tail using Accu-Check test strips (PZN 6114963, Accu Check Aviva). All mice received seven intraperitoneal injections of BrdU (50mg/kg, Sigma) at 12 months of age and were killed 28 days later with a mixture of ketamine/xylazine and a transcardial perfusion with 0.9% saline and 4% paraformaldehyde (PFA). Final blood samples were taken from the heart (right atrium) just before transcardial perfusion. Brains were left in 4% PFA over night at 4°C and were transferred to 30% sucrose before sectioning on a freezing microtome at 40 μm thickness.

### Analysis of RFID data

Radio-frequency identification (RFID) antennas, located in connecting tunnels, were used for longitudinal monitoring of behavioral activity within the cage system (Fig. 1A). Contact of mice to RFID antennas were recorded using the software PhenoSoft Control (PhenoSys GmbH, Berlin, Germany), which saved antenna and mouse identifiers together with the time stamp of the antenna contact into a database. In total 116 days/nights were recorded, however data between the 29^th^ and 42^th^, 44^th^ and 49^th^ as well as 77^th^ and 98^th^ needed to be excluded due to data loss. Raw data of the antenna contacts reduced into 5s intervals and additional meta data can be found at [data will be made available on DRYAD and link be provided at this point]. Noise reduction and calculation of RE were performed as previously described (11, 13). Because mice are nocturnal animals, only the events recorded during the dark phase were retained. Frequencies of antenna in this time series were converted to probabilities *ρ_i,j,t_* of a mouse *i* being at an antenna *j* at a night *t*. Shannon entropy of the roaming distribution was calculated as 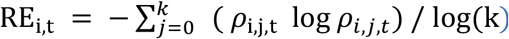, where *k* is the number of antenna. Dividing the entropy by log(k) scales the RE to the range between zero and one. Data from nights following events that could disturb patterns of exploration, such as cleaning of the cage or behavioral testing, were excluded. Cumulative RE was calculated by cumulative addition of mean RE from the 8 time blocks.

### Novel object location and recognition task

The open field and object exploration tests were performed similarly as previously described (13). Briefly, mice were placed into a square arena (40 cm × 40 cm), and their exploration was recorded using a camera (Logitech) and EthoVision software (Noldus). In total, 4 trials were performed on 3 consecutive days with a trial length of 5 min for habituation and each exploration (Fig. 4A). In the first trial (open field test), mice were put in the empty arena, which also served as habituation for the object exploration tests. In the following trial, mice were presented with 2 identical objects. The following day, one object was replaced with a “new” object. To avoid preference for object properties or placement, the position of the new object was randomized. Object A was a T25 cell culture flask with a yellow lid filled with sand [5 × 2.5 cm and 9 cm high] and object B were 3 Lego Duplo bricks in green and blue stacked to a pyramid [6.3 × 3.1 cm and 8.1 cm high]. Between each trial the arena was cleaned with 70% ethanol. Data analysis of open field and object exploration tests was performed with EthoVision as previously described (20). Briefly, the Discrimination index (DI) was calculated for trial 3 on the basis of the exploration time for the new and old object as follows: DI = (new object – old object)/(new object + old object), and ranged from −1 (preference for the old object) to 1 (preference for the new object), while 0 indicated no preference (21). For novel object recognition (NOR) at 6 month of age, object A was a plastic duck [ø 5 cm, 4.5 cm high] and B a glass bottle filled with blue liquid [ø 5cm, 10cm high]. At 6 months of age, the NOR task was followed 24 h later by the object location task, when one of the yellow ducks was displaced to the opposing corner of the area for 5 min. The time of exploration was analyzed again by DI, measuring the preference for the displaced object.

### Analysis of blood samples and ELISA Protein detection

Blood was collected into EDTA-coated microvettes (Sarstedt). After 30min to 1h, blood samples were centrifuges for 15 min at 2000 × g at room temperature. Plasma was collected from the supernatant, aliquoted and stored at −80°C. Plasma samples were used to analyze cholesterol (Amplex red cholesterol assay kit, A12216, Invitrogen), triglycerides (Triglycerides colorimetric quantification kit, ab65336, Abcam) and ApoE (mouse Apolipoprotein E assay kit, ab215086, Abcam) following the manufacturer’s instruction.

### Tissue preparation and immunohistochemistry

Tissue fixation and immunohistochemistry for the analysis of adult neurogenesis were performed as previously described (Ehret et al., 2020). Briefly, brains were cut into 40 μm coronal sections using a dry ice–cooled copper block on a sliding microtome (Leica, SM2000R). Sections were stored at 4°C in cryoprotectant solution (25% ethylene glycol and 25% glycerol in 0.1 M phosphate buffer).

For the detection of plaques, every 12^th^ section of the brain was incubated in X-34 (dissolved in 60% PBS and 40% ethanol) for 20 min, followed by 3 quick washes in tap water and development in NaOH buffer (0.2 g% NaOH in 80% ethanol) for 2min. Subsequent sections were again washed in tap water for 10 min and transferred to PBS followed by counterstaining with 5 mM Draq5 (1:500, 65-0880-92, eBiosciences) for 1h before sections were mounted in 0.1 M phosphate buffer and cover-slipped with Flouromount G (Invitrogen) mounting media. Slides were stored at 4°C till analysis at the fluorescence microscope (Axio Imager.M2, Zeiss) using a HXP a 20 × Plan-Apochromat lens and the following filter settings for X-34 (FS49, with EX BP 365 and EM BP 445/50) for Draq5 (FS50, with EX BP 640/30 and EM 690/50) and equipped with motorized stage. A tile scan of the entire hippocampus was performed. The total surface covered by amyloid plaques was determined using a custom-written script based on the “Analyze particle” function of Fiji (National Institutes of Health; http://fiji.sc/), after defining the hippocampus as region of interest. The total surface occupied by plaques was then reported and related to the hippocampal area of each section. The following functions were applied: z-projection, maximum filtration with radius of 2pixel, automatic thresholding using “Max. Entropy” and particle analysis with a circularity between 0.5 and 1.00. Automatic detection was verified manually by a blinded second investigator. For the detection of BrdU^+^ cells, the peroxidase method was applied. Briefly, free-floating sections were incubated in 0.6% hydrogen peroxide for 30 min to inhibit endogenous peroxidase activity. For antigen retrieval, sections were incubated in prewarmed 2.5 M hydrochloric acid for 30 min at 37°C, followed by extensive washes. Unspecific binding sites were blocked in tris-buffered saline (TBS) supplemented with 10% donkey serum (Jackson ImmunoResearch Labs) and 0.2% Triton X-100 (Carl Roth) for 1 h at room temperature. Primary antibodies were applied overnight at 4°C (monoclonal rat anti–BrdU 1:8000; ab6326, abcam). Sections were incubated in biotinylated secondary antibodies (Jackson ImmunoResearch Labs) for 2h at room temperature. Antibodies were diluted in TBS supplemented with 3% donkey serum and 0.2% Triton. Detection was performed using the Vectastain Elite ABC Reagent (9 μg/ml of each component: Vector Laboratories, Linaris) with diaminobenzidine (0.075 mg/ml; Sigma-Aldrich). All washing steps were performed in TBS. Stained sections were mounted onto glass slides, cleared with Neo-Clear (Millipore), and cover-slipped using Neo-Mount (Millipore). BrdU^+^ cells were counted on every 6^th^ section along the entire rostro-caudal axis of the dentate gyrus using a bright-field microscope (Leica DM 750).

For doublecortin^+^ (DCX) cell detection the peroxidase method was applied with a similar procedure but without an antigen retrieval step. As primary antibody rabbit anti-DCX (1:500, ab18723, Abcam) and biotinylated secondary antibody (donkey anti-rabbit,1:750, Jackson Immuno) were used. Analysis was performed similarly as described above for BrdU.

### Statistics

All experiments were carried out with the experimenter blinded regarding the experimental group. Statistical analyses were done using the statistical software Prism 9 (Graphpad) and R (R Core Team, 2014). In R, for normally distributed measures, we used Welsh’s t-test to compare means and F-test to test for equality of variance between groups. For repeated measures (longitudinal data), a linear mixed regression was performed using the lmer function from the lme4 package (22), and p-values were obtained by the likelihood ratio test of the full model against the model without the analyzed effects (reported previously in detail see 13). Brown-Forsythe test from the car package was used to compare the variances between groups. Two-way ANOVA was applied to identify effects through housing and genotype using Prism and R. Data were visualized using violin plot function in Prism and the ggplot2 package in R (23). Statistical results are plotted below each graph. For the histological analysis of BrdU and DCX as well as for behavioral analysis of OF, NORT and OLT, 2-way ANOVA was carried out using Bonferroni posthoc analysis.

## Results

### Enrichment at a pre-symptomatic age positively influences body weight, triglycerides and blood glucose levels in NL-F mice

NL are the best genetic control for NL-F as they differ only in the disease-promoting Beyreuther/Iberian mutation, but still carry the here apparently silent Swedish mutation. To rule out a potential interaction effect between enrichment and the Swedish mutation on general bodily metabolism, which hypothetically might be part of a systemic individualizing effect but had not been studied yet, we used C57BL/6J (C57) mice as an additional control group that also helped to further populate the large cage.

There were no differences between NL and C57 mice in body weight (Fig. 1C-E) as well as in blood glucose (Supplementary Fig. 1), cholesterol and ApoE levels throughout the experiment (Fig. 2). Triglyceride levels were slightly elevated in both NL and NL-F lines prior to but not after 4 months of ENR (Fig. 2C-D). Longitudinal analysis of body weight revealed that NL-F were heavier than NL and C57 mice already prior to ENR and after (Fig. 1D). Upon ENR, bodyweight of NL-F (but not STD) mice was consistently lower (Fig. 1C) as published for other ENR experiments (11, 13). An ENR-induced increase in variance of body weight was seen particularly in NL-F but also in the genetic controls (Fig. 1E).

**Figure 2.**
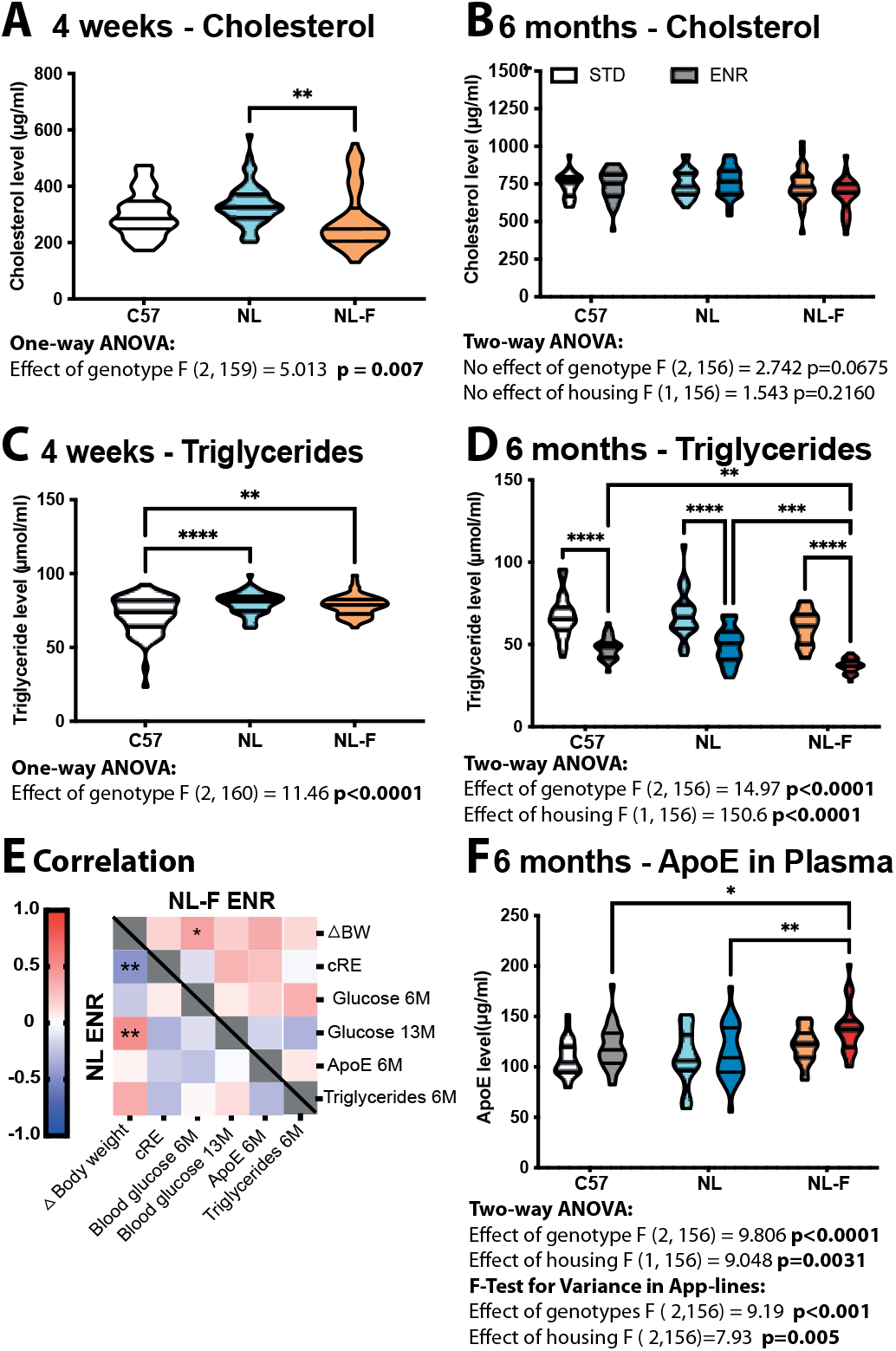
Metabolic changes in blood serum before and after complex enrichment (ENR) (**A – B**), Cholesterol levels do change upon ENR or due to AD. (**C-D**) Triglyceride levels are increased in App lines and reduce upon ENR in all lines. (**E**) Correlation of metabolic parameters and behavioural activity (cRE) and body weight are shown for both App lines in the ENR cage. A strong correlation between cumulative roaming entropy(cRE) and gain in body weight can be seen the control line (**F**) Apolipoprotein E (ApoE) is slightly elevated upon ENR housing in App NL-F/NL-F. Significant interactions are shown as *p<0.05, **p<0.01, ***p<0.005.

In line with this, an ENR-dependent reduction in blood glucose was measured only in NL-F mice in comparison to STD housed animals. Glucose levels readjusted after withdrawal from ENR but a minimal sustained effect remained significant (Supplementary Fig. 1). There is further a connection between body weight, exploration behavior and glucose levels (Fig. 2E) In addition, ENR had a positive impact on serum triglyceride levels (Fig. 2D). Interestingly, higher levels of ApoE were found in NL-F mice after ENR and additionally also a greater variance (Fig. 2F). Cholesterol levels on the other hand were not altered by ENR but decreased prior to ENR in NL-F mice (Fig. 2A-B)

Together, these results imply that ENR housing had an impact on metabolism, which is relevant in this context, because obesity, increased blood glucose and elevated triglyceride levels are risk factors of AD (1). In addition, our findings indicate that the Swedish mutation had no side effects on metabolic parameters, so that our following analyses could be based on the comparison between NL and NL-F mice as in the original publications by Saito et al. (18). Most importantly, however, we observed that ENR affected body weight trajectories and their variance in NL-F, indicating that mice with this genotype are sensitive to ENR stimuli.

### NL-F mice in long-term enriched environment show reduced inter-individual differences in behavioral trajectories

We also found that during 17 weeks of ENR NL and NL-F mice developed stable behavioral trajectories with inter-individual differences (Fig. 3A). Whereas the majority of NL animals reduced RE over time, however, NL-F animals showed a flat pattern with less variation between individuals (Fig. 3A). The analysis of changes in mean RE, as analyzed by time blocks of 2 weeks, did not significantly differ between the genotypes (data not shown). Further analysis of habituation (as a change in the slope of individual RE’s over time), however, revealed a significant reduction in habituation in NL-F (Fig. 3B). Analysis of variance, as assessed by changes in standard deviation over time, showed a reduction in total variance in NL-F (Fig. 3C). NL-F mice also showed reduced behavioral stability in comparison to controls, as detected by Pearson correlation coefficient analysis (Fig. 3D) Mice usually tend to get more predictable over time and thus Pearsons’s R increases between time blocks of analysis (13), however in NL-F this adaptation to the environment appears to be reduced. Total variance constantly increased over the entire time in the Individuality cage, so that inter-individual variance as a measure of individualization was increased after 11 weeks of enriched housing in both genotypes. Interestingly, analysis with a generalized linear mixed model revealed that this effect was not (at least not to a statistically appreciable extent) due to differences in inter-individual (residual differences) but to intra-individual differences. (Fig. 3E-G, Supplementary Table 1).

**Figure 3.**
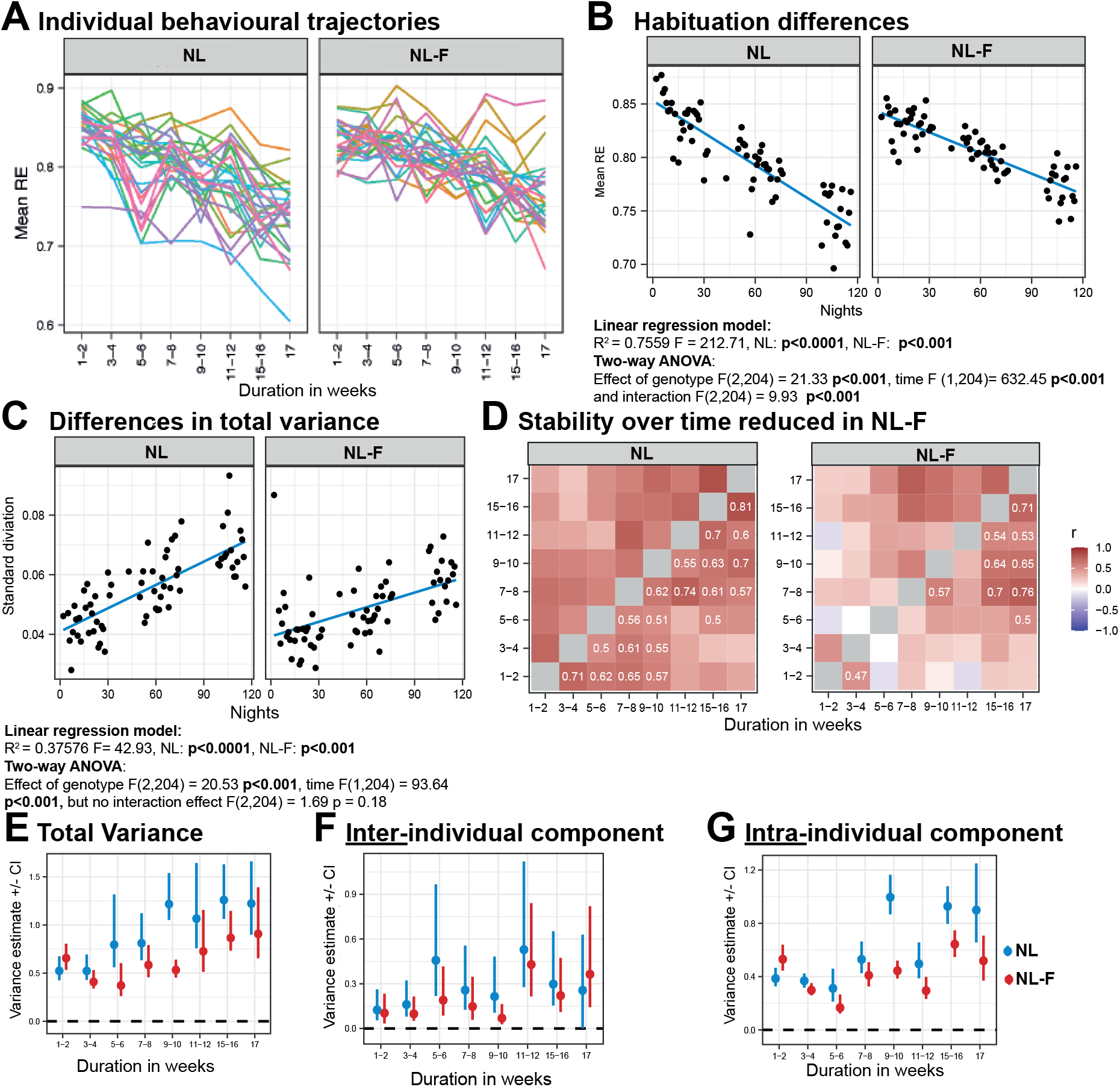
Longitudinal analysis of behavioral activity via RFID. (**A**) Individual behavioral trajectories are blotted as roaming entropy (RE) over time for the different mouse lines. (**B)** Differences in habituation are blotted as changes in mean RE over time. Linear regression analysis indicates the reduction in slope in NL-F animals. (**C**) NL-F animals have a lower variance than controls blotted by changes in standard deviation over time. (**D**) The development of stable behavioral trajectories was diminished in NL-F animals, indicating that they are less predictable. (**E-G**) Analysis of the variance component by a generalized linear mixed model (GLMM) indicates that variance between the animals from the different lines only differs in their intra-individual component (**G**) and not in their interindividual component (**F**), thus also AD animals develop individual differences over time.

**Figure 4.**
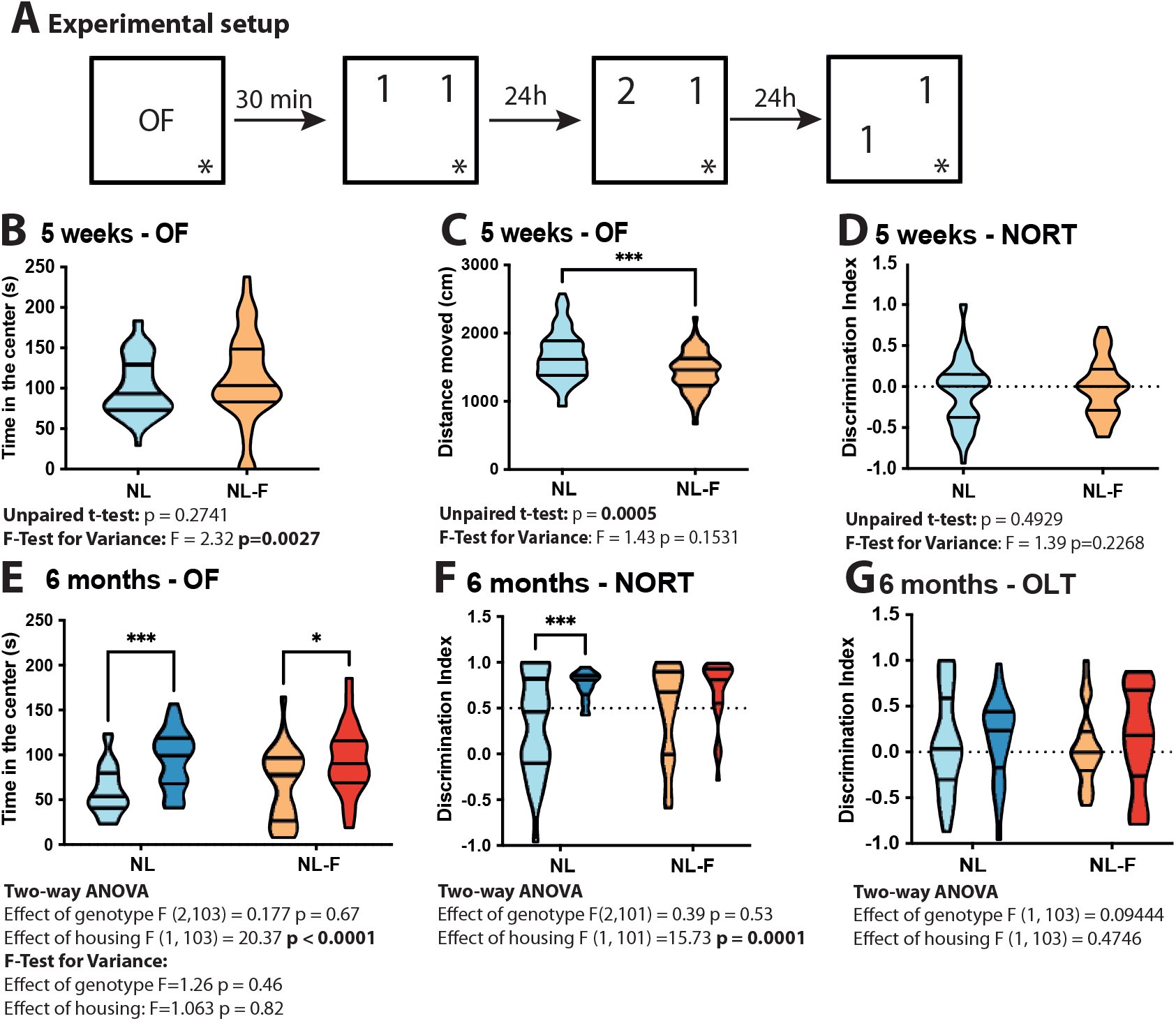
ENR increases object recognition and reduces anxiety. (**A**) Experimental setting and object placement for novel object recognition (NOR) test and object location task (OLT). (**C**) Open field activity is reduced in NL-F mice due to increased body weight. **(B, E**) After ENR, center time is increased as a measure of reduced anxiety. (**D, F**) ENR increases object recognition during NORT in NL but not NL-F. (**G**) Object displacement seems to be not improved upon ENR. Significant interactions are shown as *p<0.05, ***p<0.005.

A statistical model that contained interaction between genotype and configuration (time blocks) as a fixed term resulted in the lowest deviance information criteria (DIC) and thus reflected the underlaying data best. The inclusion of line-specific variance estimates did not improve the model when variance components were only allowed to differ along time but not between lines (Δ DIC = 2.9 only). Thus, at this pre-manifestation stage both lines showed inter-individual variation, meaning both genotypes still had a similar ability to develop individuality in terms of the means of behavioral measures. However, the intra-individual variance over time was reduced in NL-F (Fig. 3G), indicating that in NL-F individuals, we see less flexibility and less variation over time.

In conclusion, NL-F mice at this early age already showed prominent behavioral differences in longitudinal behavioral patterns. While their behavioral trajectories still had the signs of individualization, they lacked the physiological habituation and individually had become more rigid.

### ENR does not enhance object recognition in NL-F mice as in NL controls

We next asked, if there were already subtle other behavioral changes in NL-F at this early stage. We focused on open field (OF) and especially the novel object recognition task (NORT), which had shown signs of individualization in previous studies (11, 13). Prior to the exposure to ENR, there were no behavioral differences between NL and NL-F (Fig. 4B and D), except that activity of NL-F was slightly reduced (Fig. 4C). After ENR, the time spent in center of the arena was increased in the ENR groups independent of their genotype (Fig. 4E). NL mice showed an ENR-induced improvement in objection recognition (Figure 4F), but recognition of the displaced object in the object location task (OLT) was not affected (Fig. 4G). Thus, NL-F mice in ENR show a lower induction of novel object recognition compared to NL mice.

In terms of effects on individuality, we found that NL-F started out with an increased variance in the OF prior to ENR (Fig. 4 C and D), which was lost at the end of the enrichment period. At 6 months, the ENR-induced reduction in variance of the discrimination index (Fig. 4 F) was not seen in NL-F. These findings are not straightforward to interpret in line with our previous results on the individualization of OF and NORT (13, 20), but in NL-F there might be subtle very early changes in the variance of exploratory behaviors and not only their mean.

### ENR does not influence plaque load but increased variance in plaques number in the hippocampus at pre-symptomatic stage

Whether ENR has an impact on amyloid plaque load and size is still under debate and depends on the model investigated and the timing of the intervention (24, 25). We used X34 as a derivate of Congo Red for the evaluation of the plaque load (Figure 5A). Plaque load was still very low at this pre-symptomatic stage of 13 months and an ENR effect on plaque burden of the hippocampus (Fig. 5B) and cortex (Fig. 5D) was not detected, as assed by the relative area covered by plaques. Nevertheless, we revealed an ENR-induced increase in variance in plaque number (Fig. 5C), which speaks for an individualization effect on the developing AD pathology.

**Figure 5.**
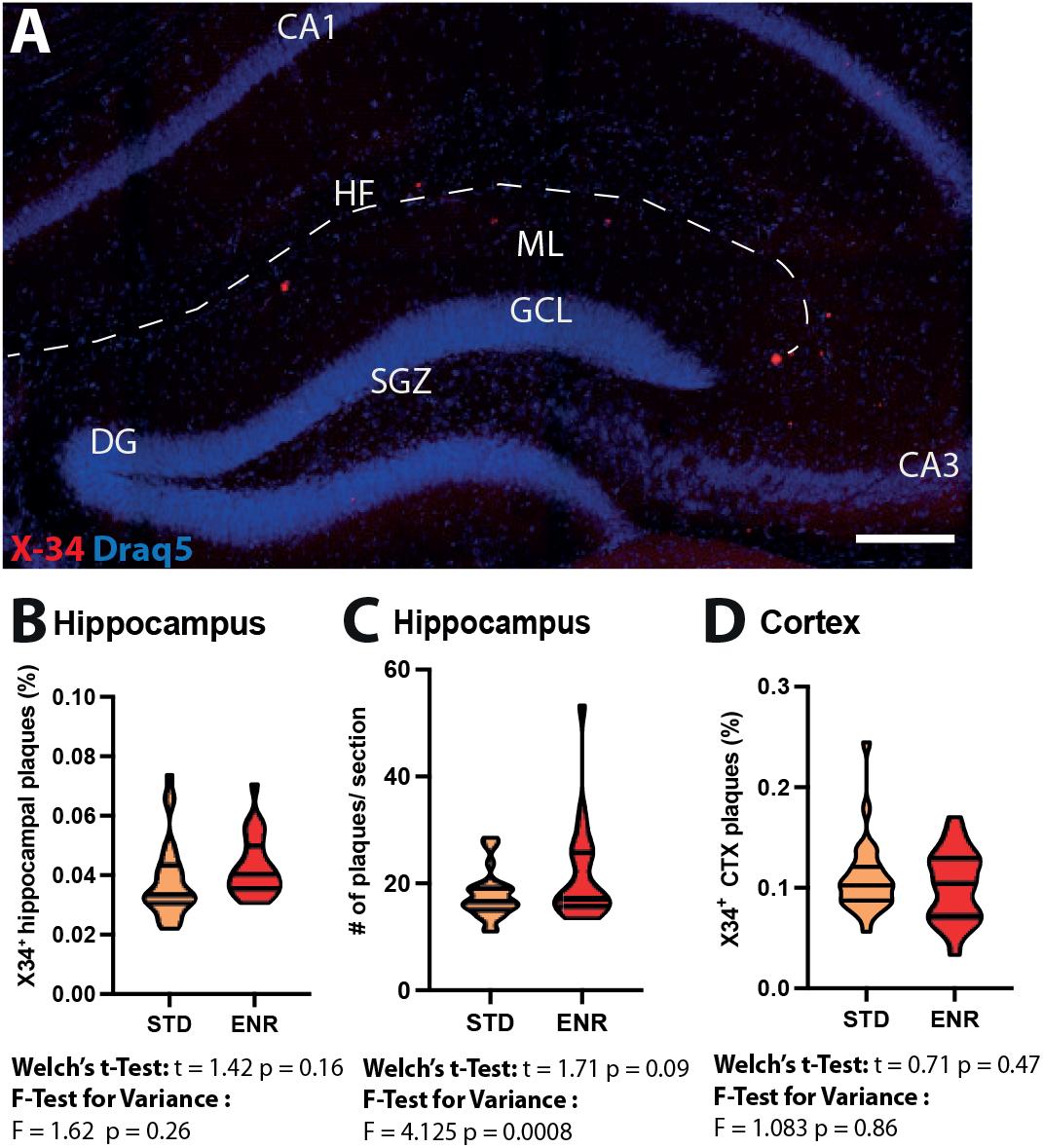
AD plaque load at an early pre-symptomatic stage of 13 months. Plaque load was analyzed by quantification of X-34 a derivate of Congo Red. (A) Representative micrograph of plaque load in the hippocampus, where plaques are shown in red. Scale bar 150 μm. Plaque coverage in hippocampus is not altered at this early stage (**B**), but an increased variance was detected by analyzing the number of plaques in hippocampus after ENR (**C**). (D) A higher plaque load can be seen in the cortex; however, no influence by ENR was detected.

### Increased adult hippopcampal neurogenesis in pre-symptomatic NL-F mice

Even though AD in human and AD-like mouse models is associated with a decrease of adult neurogenesis in late stages (16, 26–28), little is known about effects at pre-clinical stage since most animal models progress quickly and human data are scarce. The NL-F knock-in allowed us to analyze adult neurogenesis in the hippocampus at this early stage and indeed revealed an increased number of BrdU^+^ cells in the granular layer of the dentate gyrus at 28 days after the BrdU injections (Fig. 6A). To further confirm this observation with an endogenous marker, we evaluated the number and morphology of Doublecortin+ (DCX) cells (Fig. 6D). In NL-F mice, the total number of DCX^+^ cells were increased (Fig. 6B) with no significant alteration in the percentage of proliferative, intermediate or postmitotic cells (Fig. 6C). Thus, the proliferative pool was increased to the same extent as the number of postmitotic cells (Fig. 6E and F). This is consistent with the previous observation that the dynamics of maturation in adult neurogenesis is relatively stereotypic (29). The temporary increase in adult neurogenesis might reflect a compensatory response, seen also in humans (15), other mouse models (30, 31) and zebrafish (32, 33).

**Figure 6.**
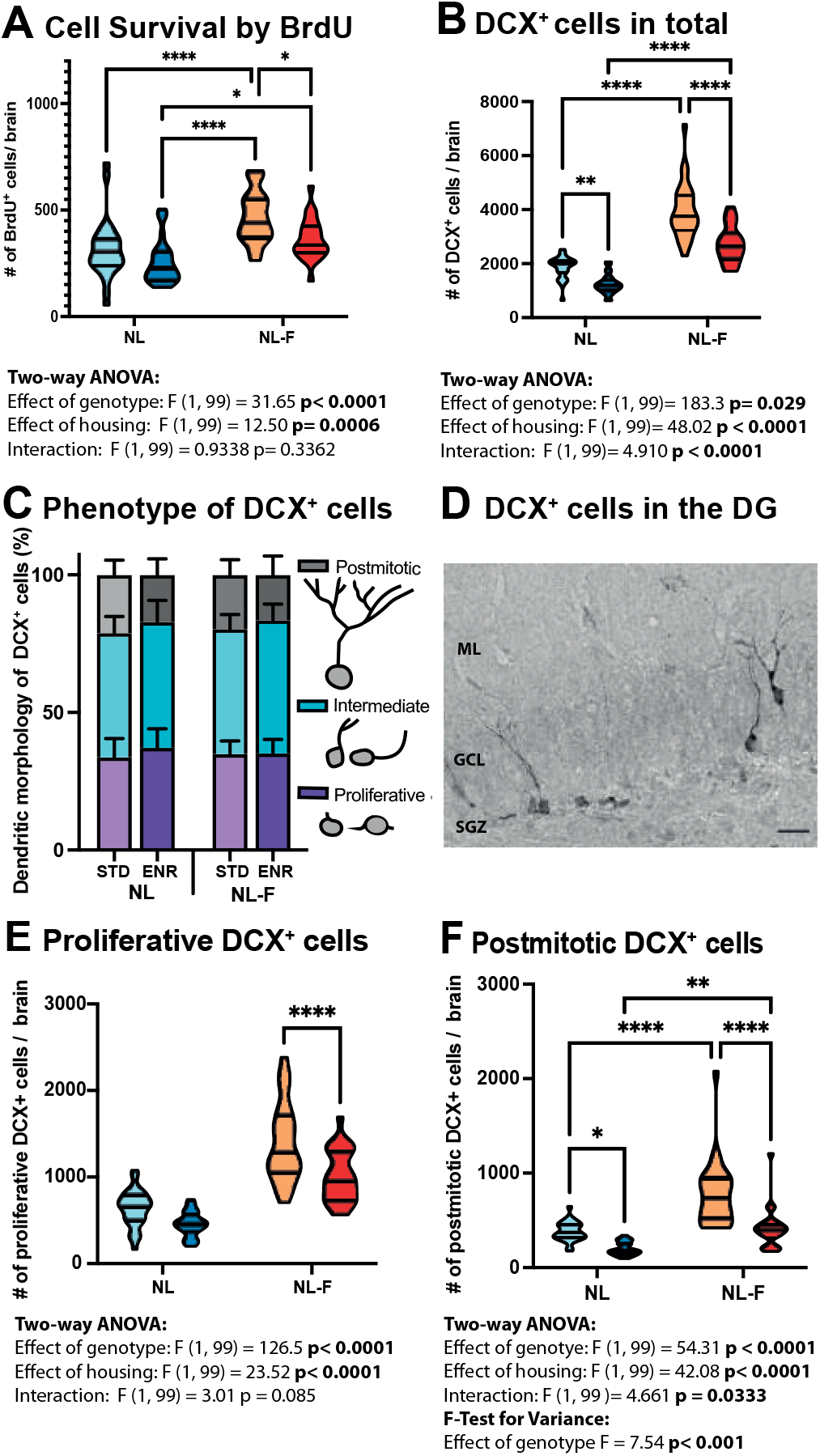
Adult neurogenesis in the hippocampus is increased at a pre-symptomatic stage in App NL-F but reduced by environmental enrichment. (**A**) The rate of adult neurogenesis in the hippocampus was assessed by BrdU incorporation in dividing cells 28 days prior to perfusion. Adult neurogenesis was increased in NL-F at this early stage and reduced in animals withdrawn from ENR. (**B**) The total number of DCX^+^ cells were increased in NL-F mice and reduced in the ENR group, similar to the BrdU quantification (**C**) Doublecortin (DCX)^+^ cells can be further classified into proliferative stage, intermediate and postmitotic stage, neither ENR nor the genotype had an effect on the percentage of differentiation. (**D**) Representative micrograph of the DCX+ cells in the dentate gyrus (DG). (**E**) The number of proliferative DCX^+^ cells after ENR withdrawal was not altered in control only in NL-F mice. (**F**) The total number of postmitotic DCX^+^ neurons were influenced by housing and genotype leading to more cells in NL-F and a compensation by ENR. Significant interactions are shown as *p<0.05 **p<0.01, ***p<0.005. Scale bar in D 25 μm.

Early exposure to ENR on the other hand reduced adult neurogenesis in the hippocampus at 12 months of age, 6 months after withdrawal from ENR. Hence, due to reduced activity and social interaction, neurogenesis was regulated back to STD levels or even slightly below in the AD condition. The well-described positive neurogenic effects of ENR only during adolescence and early adulthood is not sustained into old ages if the stimulation is not reinforced.

## Discussion

Understanding the biological mechanisms underlying individuality becomes increasingly important for personalized medicine. This is especially true in respect to dementia, because the risk of neurodegenerative diseases has a strong behavior- and lifestyle-dependent component, which might offer greater realistic benefit than so far elusive therapies. Enrichment can be considered not only as an external source of rich stimuli but also to provide the room for individual behaviors that shape the individual patterns of brain plasticity which define individuality and personality (5). This study was the first of its kind to analyze individual differences in an AD model.

We established that in a knock-in mouse model of AD complex experience in early life led to reduced inter-individual differences in behavior. The effect of early-life ENR on variance as a measure of individualization that we reported previously for wildtype mice (13), was diminished in NL-F mice. These mice became both less predictable and flexible and showed compromised object recognition even at this otherwise pre-symptomatic stage. This suggests that AD pathology very early impairs physiological individualization. In contrast, the earliest signs of plaque pathology actually showed greater variance in NL-F under ENR than under STD conditions. In addition and at first somewhat counterintuitively, after withdrawal from ENR, the pathologically enhanced adult neurogenesis that has been reported for several AD models was reduced back to normal and even slightly lower in NL-F mice. NL-F mice showed other positive responses to ENR, too: ENR positively influenced triglyceride and glucose levels in plasma.

The alteration in behavioral activity stay in close to an unpublished study comparing ENR effects in young vs old animals. There we detected more stable behavioral patterns and reduced intra-individual variance in the older animals (bioRxiv 2022.03.25.485806; doi: https://doi.org/10.1101/2022.03.25.485806)(34), suggesting that reduced adaptability is part of a reduced flexibility in older age. This remains to be confirmed, but young NL-F mice might already show behavioral alterations normally seen in older animals and thus show signs of premature aging. These mice acted less predictably. Incidentally, the feeling of “getting lost” and becoming disoriented in place and time are characteristic for AD patients (35). Hence, the observed behavioral phenotype might correspond to a very first sign of AD manifestation, long before overt plaque pathology and the typical signs of cognitive impairment.

Adult hippocampal neurogenesis, which drives the recruitment of new neurons into the existing network, might be especially important for brain maintenance, because of the strategic position of the hippocampus in complex learning and memory processes, including autobiographic memory. We have hypothesized the existence of a neurogenic reserve to cope with aging and to some degree also degenerative changes (19, 36).

The increase in adult neurogenesis might be interpreted as an initial compensatory mechanism. Such response has been found in some studies on human AD tissue (14, 15) and in murine APP_Sw,Ind_ transgenic overexpression models (30, 31). At this early stage of pathology, oligomeric Aβ might directly induce adult neurogenesis (31). In our study we also found a correlation between plaque load and BrdU levels under both housing conditions (Supplementary Fig. 3).

It is still not clear, however, whether the increase in adult neurogenesis corresponds to an abortive form of regeneration. Results from zebrafish, which show much greater endogenous potential for brain regeneration than mammals, suggest that this might be the case (33). In our analyses based on DCX expression, we saw that proliferative stages were affected similarly to postmitotic stages, so that the increase in proliferation appears to be associated with an increase in maturing postmitotic neurons (37). Whether neuronal integration at the later stages might be altered or positively influences by early ENR, remains to be studied, but we did not see any ectopic immature neurons (26) as discussed for some AD models and well-described for the increased proliferation in models of epilepsy (38, 39). Additional data indicated that trends towards a diminished number of postmitotic neurons are seen at 18 months of age (Supplementary Fig. 2). Whether there is an eventual exhaustion of the stem cell pool at later stages that might be related to the early increase in proliferation also needs to be addressed in further studies.

Over the past decades, there has been increasing evidence that changes in metabolism contribute to the development and progression of AD (40, 41) and to related risk factors like obesity. To identify whether changes in metabolism manifest already at a preclinical stage in our AD model, we investigated metabolic parameters like overall body weight, blood glucose, cholesterol and triglycerides before and after ENR. The observed increase in body weight in AD animals is in line with a previous report (41). Further, higher blood glucose levels were measured in NL-F mice in comparison the controls under STD housing conditions which reflects the previously reported alterations in glucose metabolism (42, 43) in AD patients. Since brain glucose hypometabolism precedes the development of cognitive decline in AD patients (44), an association to higher plasma glucose levels seems likely and were found previously in AD patients (45). While glucose uptake, metabolism and body weight are interconnected an AD dependent effect on glucose metabolism can be seen in this animal model already during pre-symptomatic stages of the disease.

ENR significantly counteracted the increased body weight in NL-F mice to the level of the control mice. Moreover, even after leaving the ENR, the NL-F retained a significantly lower mean blood glucose level, pointing towards a lasting effect on glucose metabolism. Increased triglyceride levels can be found in conditions like obesity, stroke and cardiovascular disease (46). ENR consistently lowered triglyceride levels across all genotypes. As these are all associated risk factors of AD, a decrease in triglycerides seem to be beneficial (47). In addition, ApoE plays a pivotal role in lipoprotein metabolism and APOE genotype has a strong impact on AD development (48), nevertheless some controversy exists in respect to ApoE levels in plasma and the risk for dementia or cardiovascular diseases (49, 50). In our study, ENR increased ApoE levels in plasma in NL-F in comparison to NL. Since triglyceride and cholesterol levels and ApoE are interconnected (51), ENR most likely has a major impact on lipid metabolism particularly in AD due to the known interactions between lipid metabolism and AD pathogenic mechanisms (reviewed by52). Astonishingly we found a negative correlation between ApoE levels and plaque area in the hippocampus of mice in ENR but not STD (Supplementary Fig. 3). Hence, in combination with the increased variance found in NL-F, individual differences in activity has a positive impact on metabolism and might also influence ApoE metabolism with potential impact on Amyloid-ß deposition and plaque burden. Thus, ENR might be able to alleviate symptoms related to metabolism in preclinical AD and thus potentially reduce the impact of related risk factors.

In conclusion, ENR is effective in NL-F but did not induce the physiological stable behavioral trajectories, long before disease manifestation. The observation that the reduced individuality was mostly explainable by the intra-individual component of the effect points to an interesting aspect that to date had received little attention. On a population scale our studies indicate that variance matters and differences might exist that lie not in the means but in the variance of these means (compare 20). We now show that something similar is true for the individual itself. Individuality also lies in behavioral flexibility and change along a more or less stable trajectory, not only in differences between those trajectories. This can be compared to the fact that information can be encoded in both frequency and amplitude of a signal. We assume (from Fig. 3F) that there are in fact also effects on the inter-individual component, which seems plausible from the appearance of the plotted trajectories in Fig. 3A, but our study turned out to be underpowered to confirm such differences. This observation underscores the necessity to gain more knowledge about what actually constitutes individuality in a neurobiological sense. Even at the most basic level of variability and variance our understanding is rudimentary, especially with respect to the contribution of intra-individual variation. A common definition of individuality (especially in the sense of personality) is one of a stable trait, but in disease contexts ‘stable instability’ might acquire a personality-like status.

Our study highlights alterations visible already in the preclinical stage of AD and emphasize the relevance of supporting innate compensatory mechanisms. In addition, this analysis strengthens the impact of enriched environment as a stimulator of metabolism and the development of individual differences in models of AD. On the other hand, constant stimulation seems necessary for direct beneficial effects of an active lifestyle on brain plasticity. Therefore, being active solely during early life is not sufficient to impact AD manifestation but fosters sustained individual differences.

In other words, while NL-F respond to the behavior-inducing environmental intervention, i.e. the shared environment, the non-shared component appears to show a reduced effect. This could mean that AD pathology, even at pre-clinical stages counteracts behavior that would support the successful build-up of resilience. While the knock-in model does not allow straightforward generalization to sporadic forms of AD and to the human situation, there is a chance of a self-reinforcing pathogenic mechanism in AD that by interfering with health behaviors determine the success of preventive life-style and activity-based interventions.

## Acknowledgements

We are particular grateful to Takashi Saito, RIKEN Brain Science Institute, for providing us with the APP knock-in models used in this study. F. Ehret received financial support from the Peter and Traudl Engelhorn Stiftung and G. Kempermann from the Alzheimer Forschung Initative (#19038). We thank Dr. Tomohisa Toda for providing aged tissue; Dr. J. Bogado Lopes, A. Karasinsky and S. Guenther for mouse handling, support in all animal experiments and legal applications. We are grateful for support from D. Lasse, C. Steinhauer and D. Glaeser on tissue processing and histology and all other members of the Kempermann laboratory for assistance and support. Further, we thank S. White at the Imaging Platform of the DZNE Dresden for support and assistance.

