## Supplemental Material for "Pre-symptomatic reduction of individuality in the App^NL-F^ knock-in model of Alzheimer’s disease"

### Supplementary Information:
#### Supplementary Figures:

**A** Blood glucose reduced in NL-F upon ENR

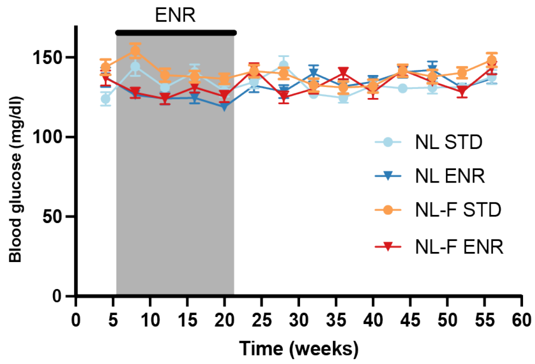


**Repeated measures ANOVA:**
F (1.859, 24.17) = 4.361 p=0.0264;
App NL-F/NL-F STD vs ENR p = 0.04

**B** No difference in glucose levels between controls

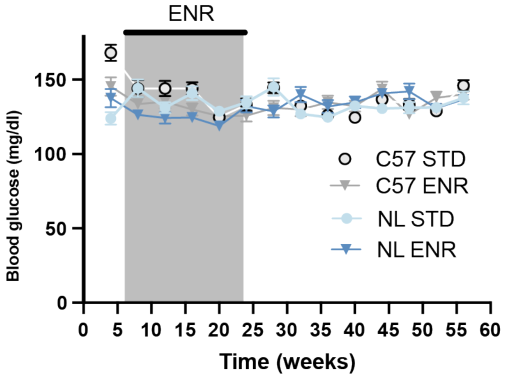


**Repeated measures ANOVA:**
F (2.428, 31.57) = 1.551 p=0.2253

**Supplementary Fig. 1: Analysis of blood glucose levels NL-F mice in comparison to the control lines C57 and NL**

(**A**)Glucose levels in NL-F are elevated in STD mice but reduce upon living in ENR in back to level of control mice. (**B**) ENR does not alter glucose metabolism in all lines only in NL-F mice.

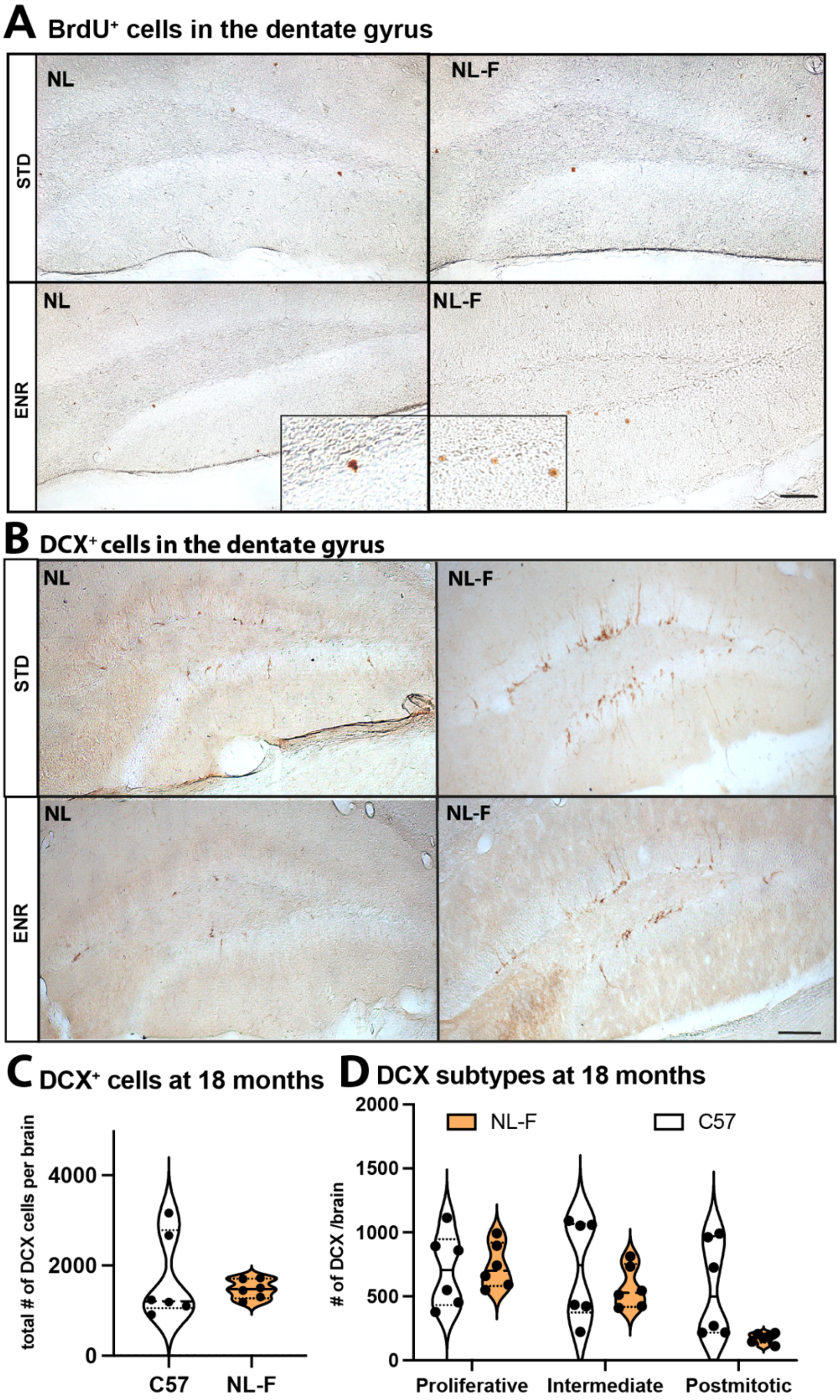


**Supplementary Fig. 2: Adult neurogenesis at 13 months and 18 months of age**

Representative images of the neurogenesis analysis by BrdU (A) and DCX (B) at 13 months of age. Analysis of Doublecortin (DCX)$^{+}$ cells in 18 months old brains from male and female NL-F and C57 controls (n = 6 per genotype, n = 3 per sex and genotype interaction). No significant changes were measured but a trend towards a decrease in the number of postmitotic cells is visible. The control mice experienced spontaneous seizures with increasing age and one mouse had enlarged ventricles and low cell numbers but due to the low number of animals available at this age we could not exclude them from the analysis. Therefore, for a more detailed analysis additional animals are required. Scale bar in A and B, 50 µm.

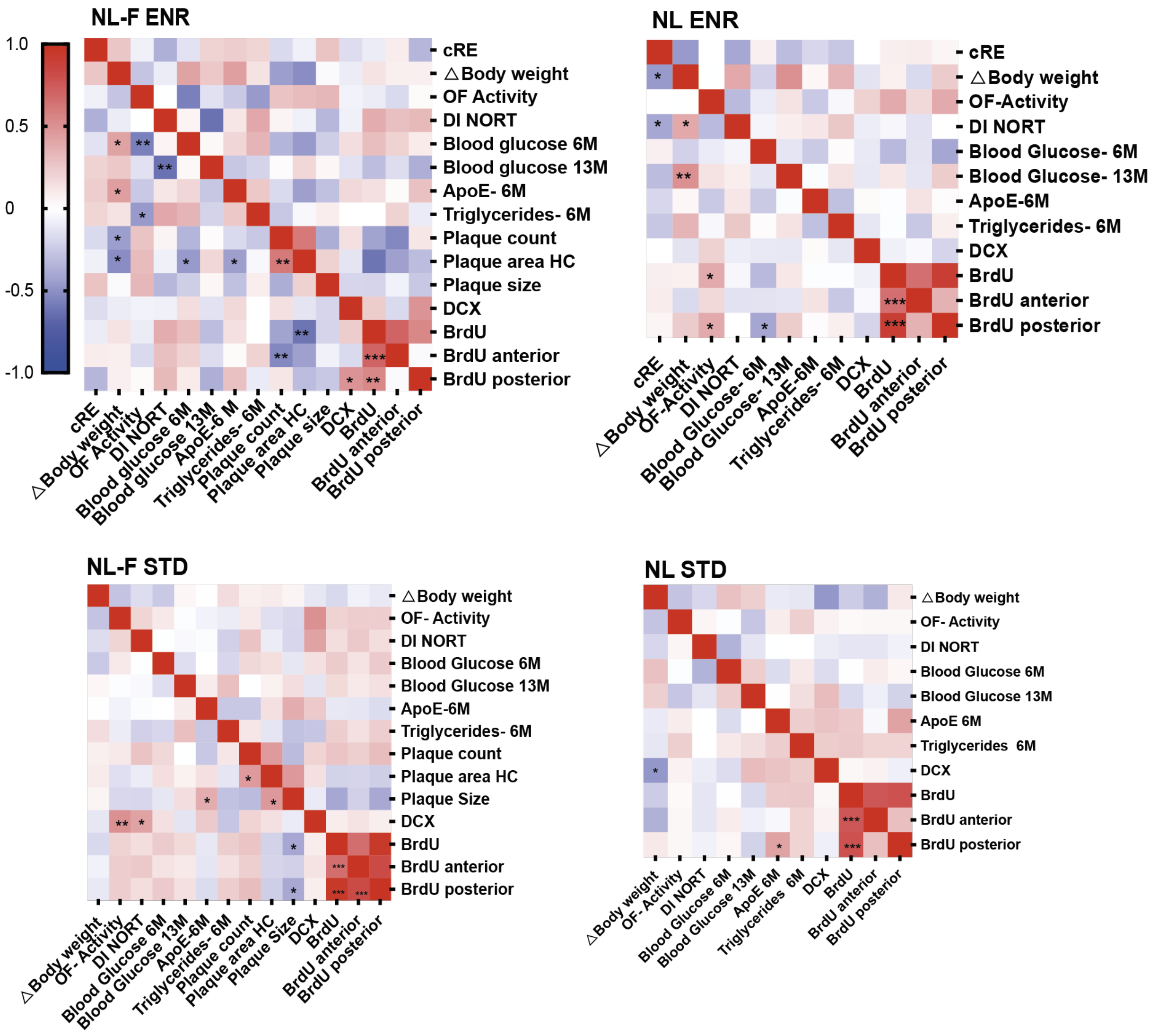


**Supplementary Fig. 3. Spearman's correlations between behavior, bodily phenotypes and adult hippocampal neurogenesis**

Heat map of Spearman's rank correlation coefficient between selected behavioral, neurogenic or metabolic traits of NL-F (left panel) and controls (NL, right panel) based on housing condition of environmental enrichment (ENR, top panels) standard (STD bottom panels). Environmental enrichment leads in NL-F mice to a negative correlation of plaque load and adult neurogenesis determined by BrdU. Further plaque area, count and size correlate. BrdU and DCX however although showing the same trend don't correlate. NL-F mice show the strongest interactions, thus parameters like body weight, blood glucose and ApoE together can give a good indication for plaque load already at a presymptomatic stage. Significant interactions are shown as *p<0.05 **p<0.01, ***p<0.005

**Supplementary Table 1: Markov Chain Monte Carlo (MCMC) analysis of a linear mixed model**

| Model | fixed.term | Inter-individual Variance | Intra-individual Variance | DIC |
|---|---|---|---|---|
| mod.int.al | line:config | line:config | line:config | 8371.49501 |
| mod.int.vres.al* | line:config | line:config | line | 8625.94805 |
| mod.int.vind.al** | line:config | line | line:config | 8783.78208 |
| mod.line.al$^{+}$ | config | config | config | 8456.77867 |
| mod.line.var.al$^{++}$ | line:config | config | config | 8453.87784 |

*Explanation: The full model (mod.int) had no intercept and contained interaction between line and config as a fixed term. Inter and intra-individual variances were estimated separately for each config and line combination, i.e. the model assumes heterogeneous inter-and intra-individual variances. To check different assumptions of the model, we calculated four models with tested terms omitted:*

*mod.int.vres - with homogenous intra-individual variance along time blocks but allowed to differ for each line

**mod.int.vind - with homogenous inter-individual variance along time blocks that was allowed to differ between lines

$^{+}$mod.line - the fixed term included only config, so this model tests whether inclusion of line improves the estimates

$^{++}$mod.line.var - the fixed term again included interaction, but both variance components were allowed to differ along time but not between lines.

*This model asks whether inclusion of interaction for variance estimation improves the estimates – or in other words whether the lines are different when it comes to development of variance over time because if this model was equally good or better than full model it would mean that variances change over time, but in the same way in all lines.*
